# Developmental plasticity in metabolism but not in energy reserve accumulation in a seasonally polyphenic butterfly

**DOI:** 10.1101/556969

**Authors:** Sami M. Kivelä, Karl Gotthard, Philipp Lehmann

**Author notes:** Corresponding author: Sami M. Kivelä.

## Abstract

The evolution of seasonal polyphenisms (discrete phenotypes in different annual generations) associated with alternative developmental pathways of diapause (overwintering) and direct development is favoured in temperate insects. Seasonal life history polyphenisms are common and include faster growth and development under direct development than diapause. However, the physiological underpinnings of this difference remain poorly known despite its significance for understanding the evolution of polyphenisms. We measured respiration and metabolic rates through the penultimate and final larval instars in the butterfly *Pieris napi* and show that directly developing larvae grew and developed faster and had a higher metabolic rate than larvae entering pupal diapause. The metabolic divergence appeared only in the final instar, that is, after the induction of developmental pathway that takes place in the penultimate instar in *P. napi*. The accumulation of fat reserves during the final larval instar was similar under diapause and direct development, which was unexpected as diapause is predicted to select for exaggerated reserve accumulation. This suggests that overwinter survival may be more dependent on metabolic suppression during diapause than the size of energy reserves. The results, nevertheless, demonstrate that physiological changes coincide with the divergence of life histories between the alternative developmental pathways, thus elucidating the proximate basis of seasonal life history polyphenisms.

**Summary statement:** The switch between diapause and direct development results in developmental pre-diapause plasticity in life history and metabolism but not in reserve accumulation in a seasonally polyphenic butterfly.

## Introduction

Temperate and high latitudes are characterized by the seasonal cycle between favourable and unfavourable conditions. Organisms living in these environments year-round must be able to tolerate adverse conditions. In short-lived organisms, survival over the adverse season is usually possible only in a specific life stage that is able to enter a state of diapause, dormancy or hibernation that can tolerate adverse conditions and survive long periods on stored energy reserves (Tauber et al., 1986; Danks, 1987; Hahn and Denlinger, 2007; Varpe, 2017). In organisms that can complete several generations during the favourable season, not all generations experience the adverse winter conditions. Hence, a developmental switch (Nijhout, 2003) resulting in alternative developmental pathways – one leading to overwintering and delayed maturation and one leading to direct maturation – evolves. Because individuals completing their life cycles during the favourable season face different selection pressures compared to overwintering individuals, the evolution of alternative developmental pathways should result in phenotypic divergence (polyphenism) between directly developing and overwintering individuals (Moran, 1992; Kivelä et al., 2013, 2016a).

Seasonal polyphenism, the expression of different phenotypes in different generations through the season, in morphology and life history is widespread in arthropods, particularly in insects (e.g. West-Eberhard, 2003; Arbačiauskas, 2004; Simpson et al., 2011). Insect seasonal life history polyphenisms are associated with alternative developmental pathways of diapause (overwintering) and direct development (e.g. Wiklund et al., 1991; Blanckenhorn and Fairbairn, 1995; Aalberg Haugen et al., 2012; Friberg et al., 2012; Välimäki et al., 2013; reviewed by Kivelä et al., 2013). Alternative life histories in the diapause and direct development pathways have a physiological basis, like life history phenotypes in general (see e.g. contributions in Flatt and Heyland, 2011). These physiological differences culminate in the life stage that is capable of entering diapause, because surviving a long adverse period in the dormant state of diapause necessitates a very different physiology compared to direct development into reproductive adult (e.g. Han and Bauce, 1998; Hahn and Denlinger, 2007, 2011; Liu et al., 2007, 2016; Rozsypal et al., 2013; Lehmann et al., 2016, 2018).

Life history divergence between the diapause and direct development pathways may, however, start accumulating already before the attainment of the developmental stage that is capable of diapause. For instance, individuals entering direct development often have shorter development times and higher growth rates compared to individuals that will enter diapause later in development (Wiklund et al., 1991; Blanckenhorn and Fairbairn, 1995; Aalberg Haugen et al., 2012; Friberg et al., 2012; Välimäki et al., 2013; reviewed by Kivelä et al., 2013). Such pre-diapause trait differentiation is expected because the developmental switch between diapause and direct development is typically triggered by environmental cues of seasonal change already before the attainment of the diapause-capable life stage (Tauber et al., 1986; Friberg et al., 2011). Hence, we would expect divergence in life history and physiology to start accumulating immediately after the induction of the developmental pathway, but the developmental timing of trait divergence between diapause and direct development remains insufficiently understood. Depending on species, life history traits start diverging between the alternative developmental pathways either early (Nylin et al., 1989; see also Lindestad et al., 2019) or late in juvenile development (Friberg et al., 2012; Esperk et al., 2013). Development and growth rates are expected to be associated with other biological rates, like metabolic rate (see e.g. Brown et al., 2004; but see Glazier, 2015 for a potential lack of association between metabolic rate and other biological rates). Nevertheless, it remains unknown whether the relatively short development time and high growth rate under direct development are associated with a high metabolic rate that also diverges after the induction of alternative developmental pathways. Alternatively, directly developing individuals may just follow the same metabolic and growth trajectories as diapausing ones but metamorphose earlier.

Besides metabolic rate, the diapause and direct development pathways are expected to show also other pre-diapause physiological differences. Overwinter survival necessitates energy reserves that can fuel metabolism through the whole dormant period during which feeding is either not possible at all or insufficient to fully fuel metabolism (Tauber et al., 1986; Danks, 1987; Hahn and Denlinger, 2007, 2011; Lehmann et al., 2016, 2018; see Liu et al., 2016 for energy reserves for summer diapause). These energy reserves must be accumulated before the beginning of diapause, which may result in differences between pathways in the allocation of resources to building reserves (Tauber et al., 1986; Danks, 1987; Hahn and Denlinger, 2007, 2011; Liu et al., 2007).

Here, we address potential pre-diapause physiological divergence between the diapause and direct development pathways by using the green-veined white butterfly *Pieris napi* (Linnaeus) as study system. *Pieris napi* produces at least two generations per year in a large part of its Palearctic distribution (Wiklund et al., 1991; Tolman and Lewington, 2002; Kivelä et al., 2015). It overwinters exclusively as a pupa, but the induction of the developmental pathway takes place during the penultimate larval instar in response to photoperiod and temperature (Friberg et al., 2011). Individuals following the diapause and direct development pathways have different life histories, including a shorter larval development time under direct development than diapause (Wiklund et al., 1991; Friberg et al., 2011; Kivelä et al., 2015). However, we do not know whether the relatively high rates of development and growth under direct development are associated with a high metabolic rate, and whether metabolic rate diverges between the diapause and direct development pathways at the same time as the divergence in development and growth rate appears. This information is vital to fully understand the physiology of alternative developmental pathways. In *P. napi*, diapause pupae contain more storage lipids than pupae that develop directly into adult six days after pupation, lipidomes and metabolomes differing between the developmental pathways in the same phase of pupal development as well (Lehmann et al., 2016, 2018). It remains unknown if these differences appear already in the larval stage during which lipid reserves are accumulated in *P. napi* (Kivelä et al., 2016b) and many other insects (Tauber et al., 1986; Hahn and Denlinger, 2007, 2011). Interestingly, however, metabolic rate and metabolome are similar in diapausing and directly developing *P. napi* pupae right after pupation (Lehmann et al., 2016, 2018). This is surprising given that the induction of the developmental pathway takes place already in the larval stage (Friberg et al., 2011) and some physiological differences appear in the final larval instar as physiology of pupation induction differs between diapause and direct development in this species (Kivelä et al., 2017). Hence, *P. napi* clearly has potential for inter-pathway physiological differentiation already in the larval stage, but whether there are differences in metabolic rate and reserve accumulation already in the larval stage needs to be elucidated.

We measure metabolic rates of growing *P. napi* larvae around the induction of either pupal diapause or direct development in the penultimate larval instar and continue the measurements through the final instar. This design enables us to assess whether metabolic rate is higher under direct development than diapause and whether potential metabolic divergence between diapause and direct development begins directly after the induction of alternative developmental pathways has taken place in the penultimate instar or later. We also measure body lipid content through the final larval instar to compare energy reserve accumulation in the alternative developmental pathways. The fat body is the major energy reserve in insects (Hahn and Denlinger, 2007, 2011; Arrese and Soulages, 2010), and since lipid is the main fuel during diapause in *P. napi* (Lehmann et al., 2016), body lipid content should be a good indicator of energy reserve size. Together, these two traits provide insight into the metabolic strategy of diapause in *P. napi*, and whether the energetic challenge is met through (i) strong metabolic suppression, (ii) extensive lipid accumulation, or (iii) a combination of both.

## Materials and methods

### Experimental design

Six diapause-generation adult *P. napi* females were captured in Stockholm, Sweden (59°22’N, 18°4’E) and individually placed in 1 L containers provided with the natural larval host plant *Alliaria petiolata* as an oviposition substrate in a laboratory. Eggs were frequently monitored and neonate larvae were individually moved to 0.5 L containers with ad libitum *A. petiolata*. Each family (offspring of a single female) was split between two climate cabinets (Termaks Series KB8400L; Termaks, Bergen, Norway): one with diapause-inducing short-day conditions (12 h day length at 20°C) and the other with long-day conditions that induce direct development (23 h day length at 20°C). The larvae were monitored daily, and fresh *A. petiolata* was provided ad libitum. As soon as the larvae started to moult to the penultimate (IV) instar, they were monitored with at most six hour intervals during the photophase and, immediately after finishing the moult, taken to the experiment. At this stage, siblings from both diapause- and direct development-inducing conditions were further split between control and respirometry groups: the first larva to finish the moult to instar IV within each family was randomized to one group, thereafter every second sibling to finish the moult being assigned to alternative groups.

Larvae in the control group were weighed (Precisa XB 120A; Precisa, Dietikon, Switzerland) twice per day with ca. 8 h interval during photophase in the short-day climate cabinet. This procedure continued through instars IV and V until the larvae became prepupae. In the respirometry group, the procedure of measuring the larvae was otherwise similar as in the control group but weighing of the larvae was always preceded by a measurement in a respirometer (larvae were weighed immediately after the respirometric measurement; see below for details of respirometric measurements). Larvae were kept in the climate cabinets under the assigned diapause- or direct development-inducing conditions all the time, except during the measurements.

Prepupae were monitored twice a day for pupation and, approximately 48 h after pupation, the pupae were weighed and sex was determined. Pupae were housed under the same conditions in the same climate cabinets where they were during the larval period. Pupae were monitored daily for adult eclosion. At 20°C, directly developing individuals typically eclose within two weeks after pupation. Therefore, we considered all individuals that did not eclose and showed no signs of development within three weeks after pupation to be in diapause. All such individuals were still in diapause after six months.

The experiment was repeated with two temporally separate cohorts so that the start of the experiment in the second cohort was 14 days later than that in the first cohort. Different families were included in the two cohorts. The first and the second cohorts included 2 and 4 families, respectively. In the first cohort, 10 control group larvae and 10 respirometry group larvae were exposed to both diapause- and direct development-inducing conditions (5 larvae/family in each combination of developmental pathway and group). In the second cohort, the group-specific number of larvae was 12 (2–5 larvae/family in each combination of developmental pathway and group). Due to mortality in the first cohort (two direct-development control larvae), we got data from 42 control and 44 respirometry larvae through the instars IV and V in total.

### Respirometry

Differential mode flow-through respirometry was used to measure carbon dioxide (CO_2_) production and oxygen (O_2_) consumption of individual larvae. Outdoor air was sucked into the system by SS-4 pump (Sable Systems, Las Vegas, USA) set to 500 ml min^−1^, and scrubbed of H_2_O using drierite (WA Hammond Drierite, Xenia, USA). Then, the air inflow was split into two lines. Both lines were pushed at a steady rate of 150 ml/min using two separate mass flow controllers (840 Series; Sierra Instruments Inc, California, USA). The first line went into a MUX multiplexer (Sable Systems) and through the measurement chamber (volume 10 ml), located in a climatic cabinet (Panasonic MIR-154-PE, Panasonic, Osaka, Japan) set to 20°C. Thereafter, it again was scrubbed of H_2_O with magnesium perchlorate (Sigma-Aldrich) before entering the sample line of a Li-7000 CO_2_ analyser (LiCor, Lincoln, USA). The second line proceeded in the same way, mimicking the exact length of the sample line (including an empty measurement chamber), before entering the reference line of the Li-7000 CO_2_ analyser. Both lines then proceeded through ascarite CO_2_ scrubbers (Acros Organics, New Jersey, USA), and entered an Oxzilla FC-2 O_2_ analyser (Sable Systems) after which air was ejected to the room. Both CO_2_ and O_2_ analysers were calibrated before the start of the experiment. Differential CO_2_ and O_2_ were calculated by subtracting the output of the reference line from the output of the sample line. Sampling rate was 1 Hz for all measurements. The measurement protocol entailed a 2 min baseline measurement of an empty chamber (preliminary experiments showed that this was long enough), followed by an 8 min measurement of the chamber containing the animal, and ending with a 3 min measurement of the empty chamber (see Fig. S1). The raw data was first lag corrected to synchronize CO_2_ and O_2_ traces and then drift corrected (there was minimal drift during the 15 minute measurements). The trace was then baseline corrected against the empty chamber value, fractioned and multiplied with flow rate to convert the raw output to ml CO_2_ or O_2_ h^−1^ (Lighton 2008).

We aimed to minimise the time larvae had to be without access to food, and the consequent negative effects on growth. Therefore, larvae were not starved before respirometric measurements, yet they were without food during the measurement (lasting approximately 15 minutes including handling). The number of faecal pellets produced during the measurement was recorded to take into account potential variation in the digestive cycle and CO_2_ released from the faeces. We did not measure movement of larvae during measurements as a previous study using a similar setup showed that larvae move little in the tubes and, in that study, the movement effect was either statistically insignificant or very small (Kivelä et al., 2016b).

We calculated RQ only from measurements where signal-to-noise ratio was >5 as these measurements were considered as reliable. The signal-to-noise ratio was calculated as the difference in measurement mean and baseline mean, divided by the baseline standard deviation. All CO_2_ measurements exceeded this threshold but, for O2, the signal was generally too weak until the end of the penultimate instar (the smallest larva with a successful O_2_ measurement was 18.4 mg).

### Analysis of body lipid and water content

In addition to larvae used in control and respirometric measurements, we reared extra larvae under the above-mentioned diapause and direct development conditions. Once these larvae entered the final (V) instar, a time point (day during the instar V or a two-day old pupa) for freezing was randomized for each individual. The larvae were weighed twice a day until they were frozen in liquid nitrogen at the pre-determined time points. The numbers of frozen final-instar larvae and pupae were 99 and 14 in the diapause pathway, respectively, the corresponding direct development pathway numbers being 94 and 17.

Samples were kept at −80°C until analysis. Water and total lipid content of larvae was measured after chloroform:methanol (Chl:Meth) extraction (Folch et al., 1957) according to the following protocol. First, larvae were weighed (fresh mass; Precisa XB 120A balance), and then dried for 72 h at 55°C. Larvae were then reweighed (dry mass), placed in small glass vials (20 ml) containing 4 ml Chl:Meth (2:1 [vol:vol]), crushed thoroughly with a glass rod, and left for 72 h in a fume hood at 20°C. Chl:Meth effectively dissolves lipids in the sample into the liquid phase. This includes neutral lipids and phospholipids, but also glycerol, carbohydrates and amino acids (Newman et al., 1972), so this method overestimates total lipids (Williams et al., 2011). In *P. napi*, there appears to be no significant accumulation of these compounds right after pupation regardless of pathway (Lehmann et al., 2018) and, therefore, the method should be suitable for between-pathway comparisons even though the absolute lipid values need to be treated with caution. A previous study, using mass-spectrometric methods, verifies that neutral lipids constitute the absolute major component of total lipids in young pupae of *P. napi* (Lehmann et al., 2016). After the Chl:Meth submersion, 250 μl of the suspension was retrieved with a syringe (Hamilton, Model 1825 RN SYR, 22s ga) and applied in pre-weighed (Cahn 28 Automatic Electrobalance), aluminium cups (350 μl VWR Aluminium round micro weigh). Between each sample the syringe was rinsed with Chl:Meth twice. The aluminium cups were left to evaporate in a fume hood for 4 hours and reweighed.

Total lipid mass in the sample was calculated by multiplying the mass of lipids in the cup with the dilution factor (4000/250 = 16). Relative lipid content was calculated as the ratio of total lipid mass to dry mass. Relative water content was calculated as 1 – ([dry mass] / [fresh mass]).

### Statistical analyses

We used linear mixed-effects models fitted with the maximum likelihood method by using the function ‘lme’ (package ‘nlme’; Pinheiro et al., 2016) in R version 3.3.2 (R Core Team, 2016) in all data analyses. The multimodel inference approach (e.g. Burnham and Anderson, 2002) was used throughout.

We compared life history traits associated with growth and development (development time through instars IV and V, growth rate in instars IV and V, time from the beginning of instar IV until the attainment of peak larval mass in instar V, peak larval mass, and pupal mass) between the diapause and direct development pathways, and between the control and respirometry groups. Individual-specific growth rate was calculated as the regression slope of ln(mass) on time (days) in the focal instar by including all observations in the regression in instar IV, and the first half of them in instar V to get a growth rate estimate for the free growth period before growth deceleration preceding pupation (see Fig. S2). In analyses of life history traits, the “global model” (i.e. the most extensive model considered) included group (control/respirometry), developmental pathway (diapause/direct), sex (female/male) and all interactions among them, as well as cohort (first/second; not included in any interactions), as fixed effects. Family was set as a random effect to control for non-independence of siblings. The set of all meaningful models (i.e. following the principle of hierarchy) simpler than the global model was derived with the function ‘dredge’ (package ‘MuMIn’; Barton, 2016; see Oline Resource, Table S1, for the sets of models with Akaike weights >0.05). The models, including the global model, were averaged with the function ‘model.avg’ (Barton, 2016). When needed, we took heteroscedasticity into account by adding weights determined by a variance function (either ‘varIdent’ or ‘varExp’ [Pinheiro et al., 2016]) into the models. The sets of models with Akaike weight >0.05 are presented in Table S1.

For CO_2_ production rate, we analysed instars IV and V separately and included only observations during the growth period in the analysis. Data were ln-transformed to facilitate allometric inferences (see Kilmer and Rodríguez, 2016 for arguments favouring this approach over reduced major axis regression). Fixed effects in the global model included centered ln-transformed mass, developmental pathway, sex and all interactions among them, as well as the square of centered ln-transformed mass included in interactions with developmental pathway and sex. Cohort (not included in any interactions) and the number of frass produced during the respirometric measurement (a covariate) were also included in the fixed effects. To control for repeated within-individual measures, random effects included random individual-specific intercepts and slopes in relation to centered ln-transformed mass. The inclusion of random slopes significantly improved model goodness-of-fit over random-intercepts-only model both in instar IV and V data when measured by Akaike information criterion (ΔAIC>2). The inclusion of random individual-specific slopes is also important to avoid overconfident estimates of fixed effects (Schielzeth and Forstmeier, 2009). To reduce correlations among random intercepts and slopes, ln-transformed mass was centered (subtracting the mean from each observation). Centering a variable only changes the mean but not the variance. Therefore, centering of ln(mass) does not affect the regression slopes of other traits on it, and allometric inferences remain, consequently, unaffected by centering. Model averaging and modelling of heteroscedasticity was performed as explained above, and the sets of models with Akaike weight >0.05 are presented in Table S2.

Variation in RQ was analysed similarly as explained above for CO_2_, except that the models included only random individual-specific intercepts because the inclusion of random individual-specific slopes did not improve model goodness-of-fit (AIC values were lower for the random intercept models). Because there was no need to include random individual-specific slopes in the models for RQ, mass was not centered to make interpretation of the results easier. In instar IV, we added weights by the ‘varIdent’ variance function (Pinheiro et al., 2016) to the models, because modelling the higher residual variance in females than in males improved the estimation accuracy of random effects, yet AIC did not suggest a difference between the weighted and unweighted models (ΔAIC= 0.919). The sets of models with Akaike weight >0.05 are presented in Table S3.

In the analyses of body composition data (body water and lipid content) from instar V larvae, fixed effects of the global model included developmental pathway, ln-transformed fresh mass, the square of ln-transformed fresh mass and interactions between developmental pathway and each of ln-transformed fresh mass and its square. Family was set as a random effect. Weights by the ‘varExp’ variance function (Pinheiro et al., 2016) significantly improved model goodness-of-fit (ΔAIC=43.0 for water content and ΔAIC=11.5 for lipid content). We also compared relative water and lipid contents between similarly-sized instar V larvae and pupae. Here, only larvae whose mass was between the minimum and maximum of pupal masses in these data were included in the analysis. The global model included developmental pathway, developmental stage (instar V larva/pupa) and their interaction as fixed effects, and family as a random effect. Model averaging and modelling of heteroscedasticity was performed as explained above, and the sets of models with Akaike weight >0.05 are presented in Tables S4 and S5.

## Results

### Life history divergence between alternative developmental pathways

In both the control and respirometry group, all individuals exposed to the short day length manipulation entered diapause, while the long day manipulation invariably induced direct development. Individuals developing directly into adults had higher larval growth rates than individuals entering pupal diapause both in instars IV and V (Table 1; Fig 1). Development time through instars IV and V was shorter under direct development than diapause, and directly developing individuals reached the peak larval mass in instar V sooner after entering instar IV than the ones entering diapause (Table 1; Fig. 2). Instar duration was shorter under direct development than diapause both in instar IV (direct development mean=2.11 [95% CI 2.03, 2.19] days; diapause mean=2.62 [2.52, 2.72] days) and V (direct development mean= 3.38 [3.19, 3.57] days; diapause mean= 4.64 [4.49, 4.79] days), the relative difference in instar durations becoming more pronounced in instar V compared to instar IV (instar duration under direct development is 73% and 81% of that under diapause development in instars V and IV, respectively). Despite having higher growth rate, the fast development of directly maturing individuals led to a low peak larval mass compared to that attained by individuals entering diapause (Table 1; Fig. 2). Males had higher pupal masses than females, and there was also a weak tendency for the pupal mass to be higher under diapause than direct development (Table 1; Fig. 2).

**Figure 1.**
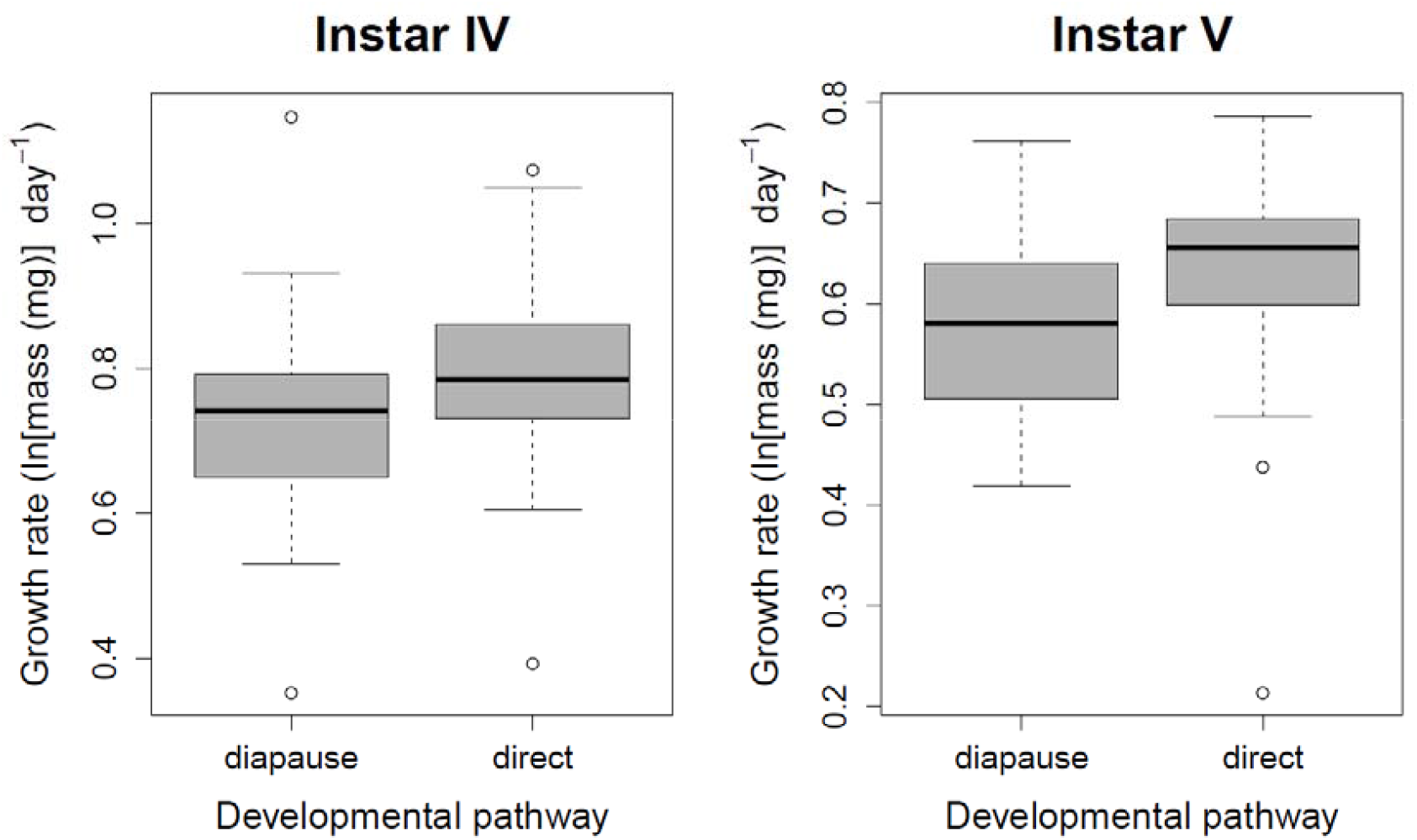
Larval growth rates under diapause and direct development in instars IV (left) and V (right). Each box includes 50% of observations, the horizontal line within the box depicting the median value. The whiskers depict minimum and maximum of the distribution while the circles depict outliers.

**Figure 2.**
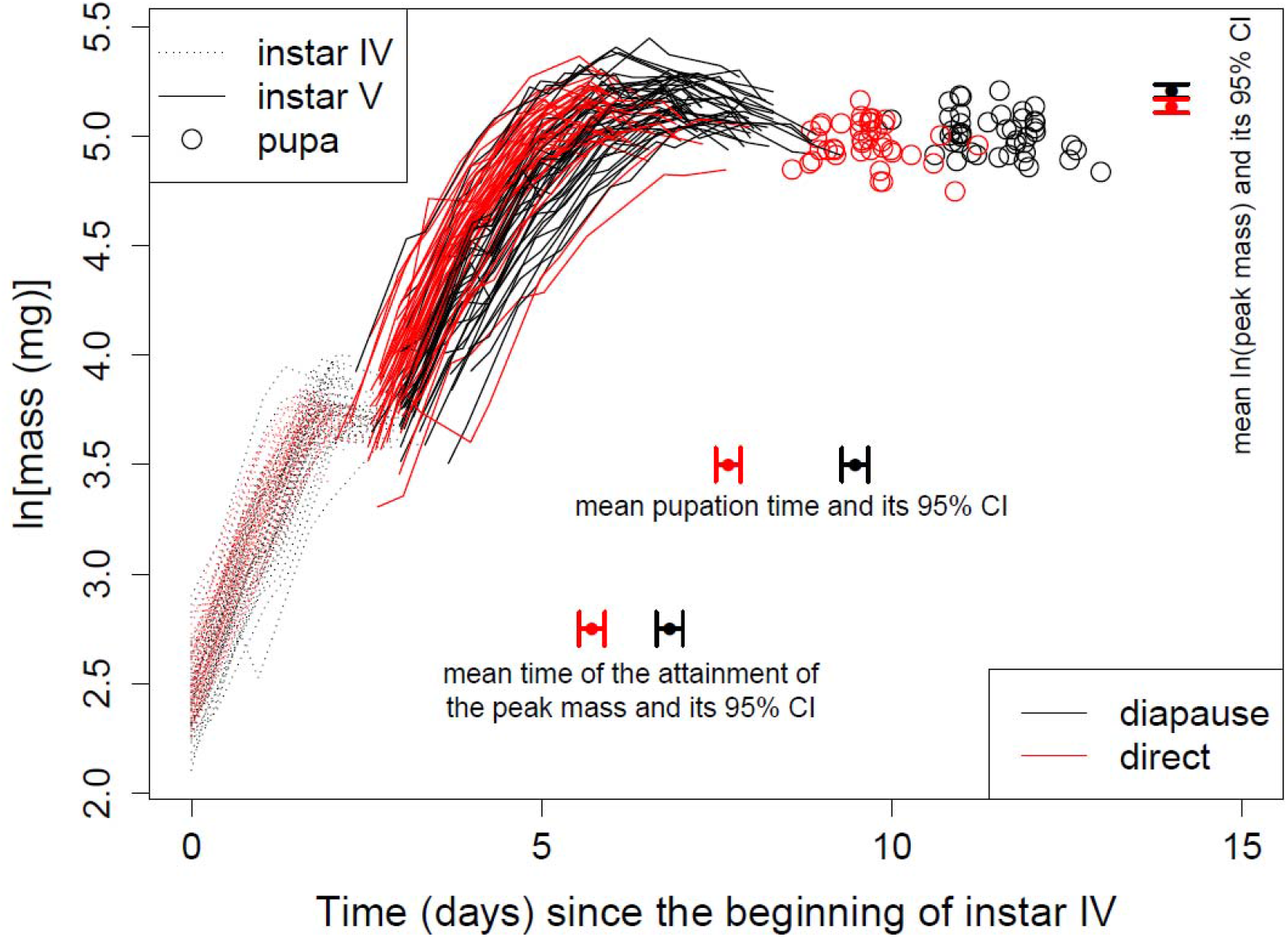
Growth trajectories of experimental larvae (both control and respirometry group larvae are included) through instars IV (dotted lines) and V (continuous lines). Black and red lines depict growth trajectories of individuals entering pupal diapause and direct development into adult, respectively. Open circles depict pupal masses two days after pupation. Closed circles below the growth trajectories indicate mean times of the attainment of peak larval mass and pupation in the diapause (black points) and direct development (red points), while horizontal whiskers illustrate their 95% confidence intervals (CI). Closed black and red circles and associated vertical whiskers at top right corner indicate mean peak larval masses (in ln-scale) and their 95% CIs under diapause and direct development, respectively.

**Table 1.** Statistically significant (risk level 0.1) model-averaged (full average) fixed effects of linear mixed-effects models fitted with the maximum likelihood method explaining life-history variation between alternative developmental pathways (diapause / direct development), experimental groups (control / respirometry) and cohorts (first / second). The models included random family-specific intercepts.

| Trait | Term (relative importance) | Factor levels | Averaged estimate | Adjusted S.E. | z-value | P-value |
| --- | --- | --- | --- | --- | --- | --- |
| Development time |  | Intercept | 9.28 | 0.192 | 48.3 | <0.0001 |
|  | Pathway (1) | direct | -1.80 | 0.149 | 12.1 | <0.0001 |
| Time until peak mass |  | Intercept | 6.72 | 0.146 | 46.0 | <0.0001 |
|  | Pathway (1) | direct | -1.11 | 0.150 | 7.38 | <0.0001 |
| Peak mass |  | Intercept | 177 | 5.35 | 33.1 | <0.0001 |
|  | Cohort (0.92) | second | 15.0 | 6.99 | 2.14 | 0.032 |
|  | Group (0.99) | respirometry | -10.1 | 4.04 | 2.50 | 0.012 |
|  | Pathway (1) | direct | -12.1 | 4.26 | 2.84 | 0.0044 |
| Pupal mass |  | Intercept | 146 | 4.16 | 35.0 | <0.0001 |
|  | Pathway (0.94) | direct | -6.22 | 3.51 | 1.78 | 0.075 |
|  | Sex (0.99) | male | 9.04 | 3.37 | 2.68 | 0.0073 |
| Growth rate IV<sup>a</sup> |  | Intercept | 0.728 | 0.0274 | 26.6 | <0.0001 |
|  | Pathway (0.98) | direct | 0.0907 | 0.0313 | 2.90 | 0.0037 |
| Growth rate V<sup>b</sup> |  | Intercept | 0.589 | 0.0233 | 25.3 | <0.0001 |
|  | Pathway (1) | direct | 0.0855 | 0.0287 | 2.98 | 0.0029 |

<sup>a</sup> 'varIdent' variance function to take different variances in different families into account.
<sup>b</sup> 'varIdent' variance function to take different variances in females and males into account.

### Metabolic divergence between alternative developmental pathways

We use whole-organism CO_2_ production rate as a proxy for metabolic rate because there is a strong and positive correlation between these two traits (estimated correlation coefficient based on resampling to avoid pseudoreplication = 0.961 [95% percentile CI 0.934, 0.980]; Fig. S3). Furthermore, O_2_ consumption rate, that is needed in the calculation of metabolic rate, could not be reliably measured early in instar IV with the set-up used here. Therefore, to facilitate inter-pathway comparisons through the two investigated instars, we base our inferences on CO_2_ production rate.

CO_2_ production rate increased non-linearly through instar IV; a hyperallometric increase early in the instar changed to a hypoallometric increase later in the instar, and the ontogenetic increase in CO_2_ production ceased altogether at the end of the instar (Table 2; Fig. 3a, c; see also Fig. S4). There was no difference between diapause and direct development in CO_2_ production in instar IV (Table 2). In instar V, CO_2_ production had a qualitatively similar relationship to body mass as in instar IV, yet the non-linearity of the relationship was weaker, and there was a difference between the alternative developmental pathways, CO_2_ production being higher under direct development than diapause (Table 2; Fig. 3b, d).

**Figure 3.**
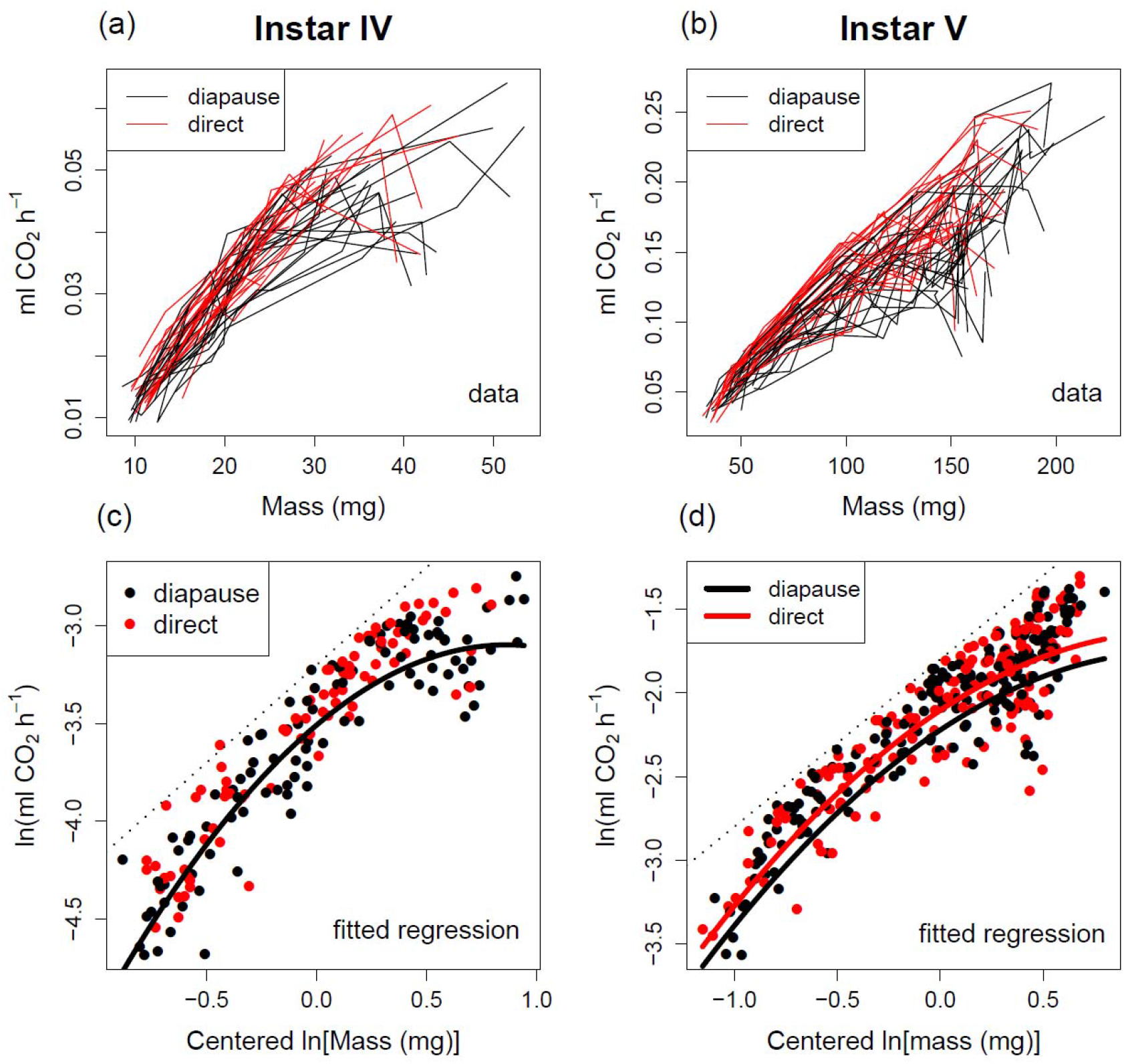
Individual-specific trajectories of changing whole-organism CO_2_ production rate (lines) along with growth (increasing mass) under diapause (black lines) and direct development (red lines) in instars IV (a) and V (b). The bottom row illustrates allometric regression curves fitted to ln-transformed data (points) in instars IV (c) and V (d). The regression curves are based on model-averaged coefficients (see Table 2) fitted to data including centered ln(mass) and illustrate within-instar allometry of CO_2_ production rate. There was no difference between diapause and direct development in instar V, so only a single regression curve is presented in (c). The dotted lines in the bottom panels are isometric reference lines with arbitrary intercepts.

**Table 2.**
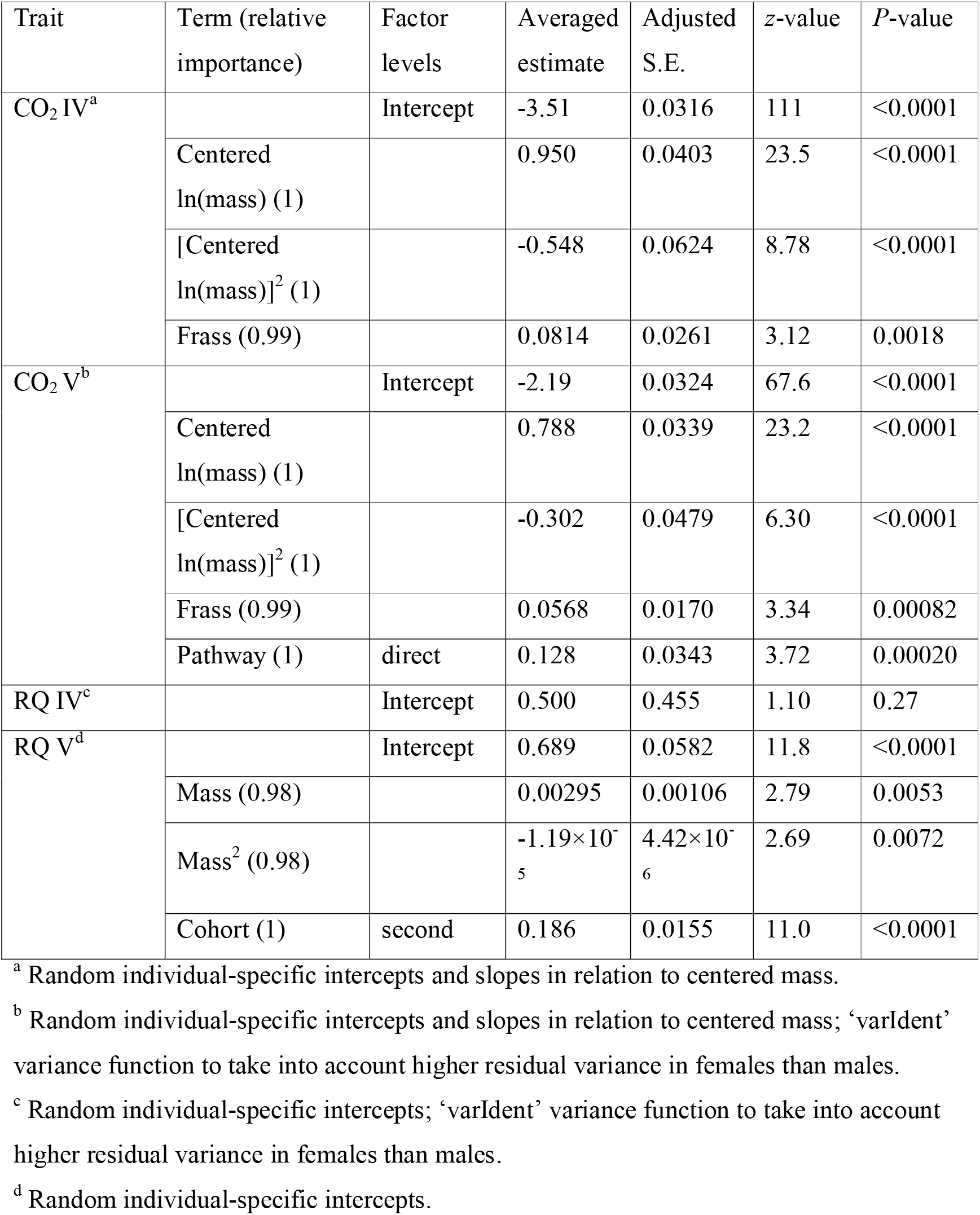
Statistically significant (risk level 0.1; insignificant lower-level terms are presented for interactions and squared effects; at least an intercept is presented for each trait) model-averaged (full average) fixed effects of linear mixed-effects models fitted with the maximum likelihood method explaining variation in CO_2_ production rate and respiratory quotient (RQ) between alternative developmental pathways (diapause / direct development) and experimental cohorts (first / second). CO_2_ production rate was ln-transformed in both instars.

The respiratory quotient (RQ) in instar IV was independent of developmental pathway and body mass, the latter being apparently due to the data concentrating at the latter part of the instar (Table 2; Fig. 4a). The larvae became large enough for reliable measurement of O_2_ consumption (and RQ) only in the latter part of the instar IV, which is why we do not have RQ data from the beginning of the instar IV. In instar V, RQ showed a peaked relationship to body mass, RQ values being higher in the second than in the first cohort (Table 2; Fig. 4b). There was no difference in RQ values between diapause and direct development.

**Figure 4.**
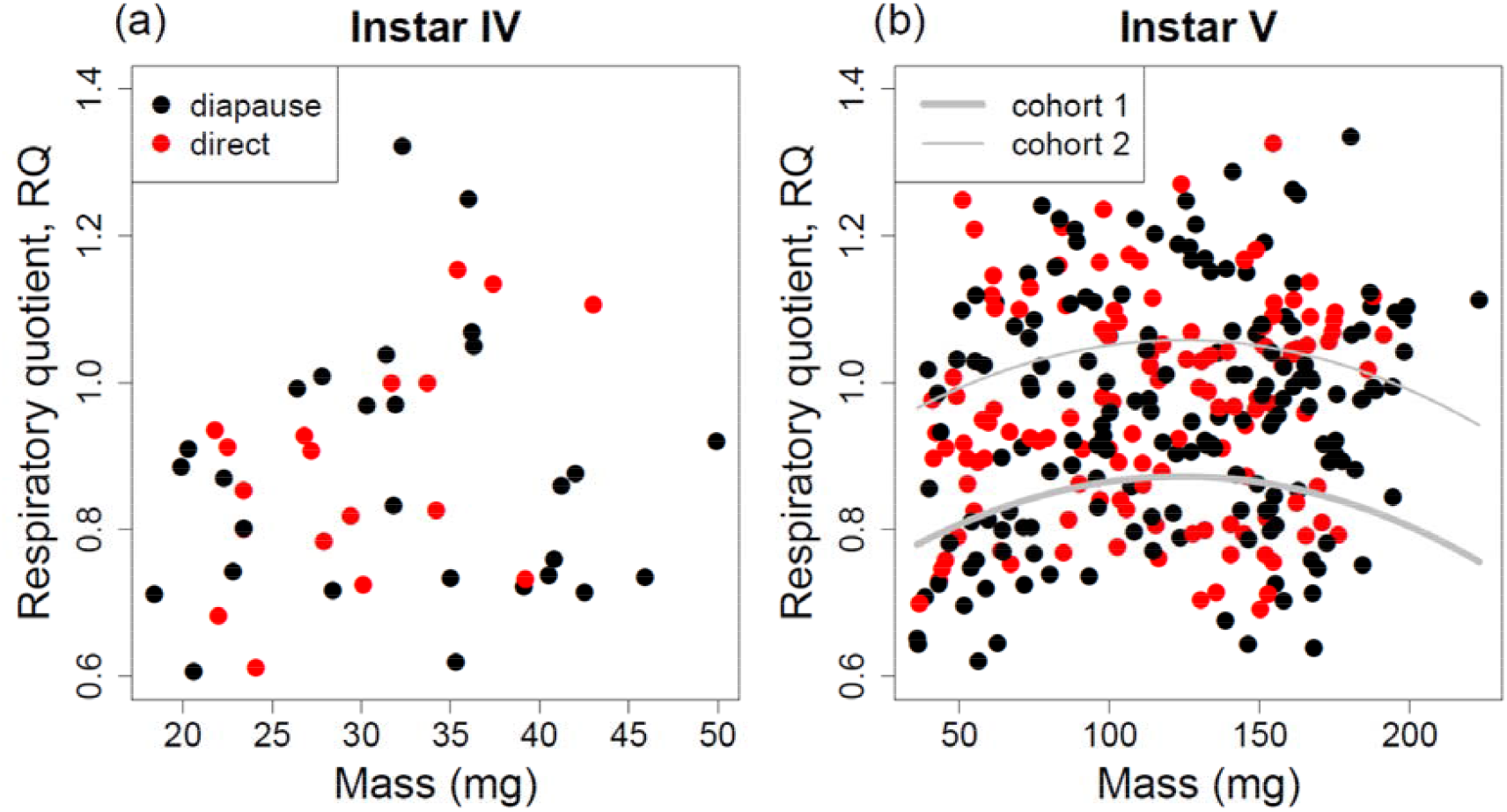
Respiratory quotient, RQ, in relation to body mass in instars IV (a) and V (b). Points depict observations. The grey curves in (b) are cohort-specific regression curves based on the model-averaged coefficients (see Table 2).

### Body water and lipid contents in alternative developmental pathways

The absolute amount of water in the body did not differ between diapause and direct development and increased nearly isometrically, yet non-linearly, over the instar V, the scaling becoming increasingly hypoallometric at the end of the instar (Table 3; Fig. 5a). Absolute mass of lipids in the body did not differ between diapause and direct development either and increased linearly and hyperallometrically through instar V (allometric exponent=1.37; Table 3; Fig. 5b).

**Figure 5.**
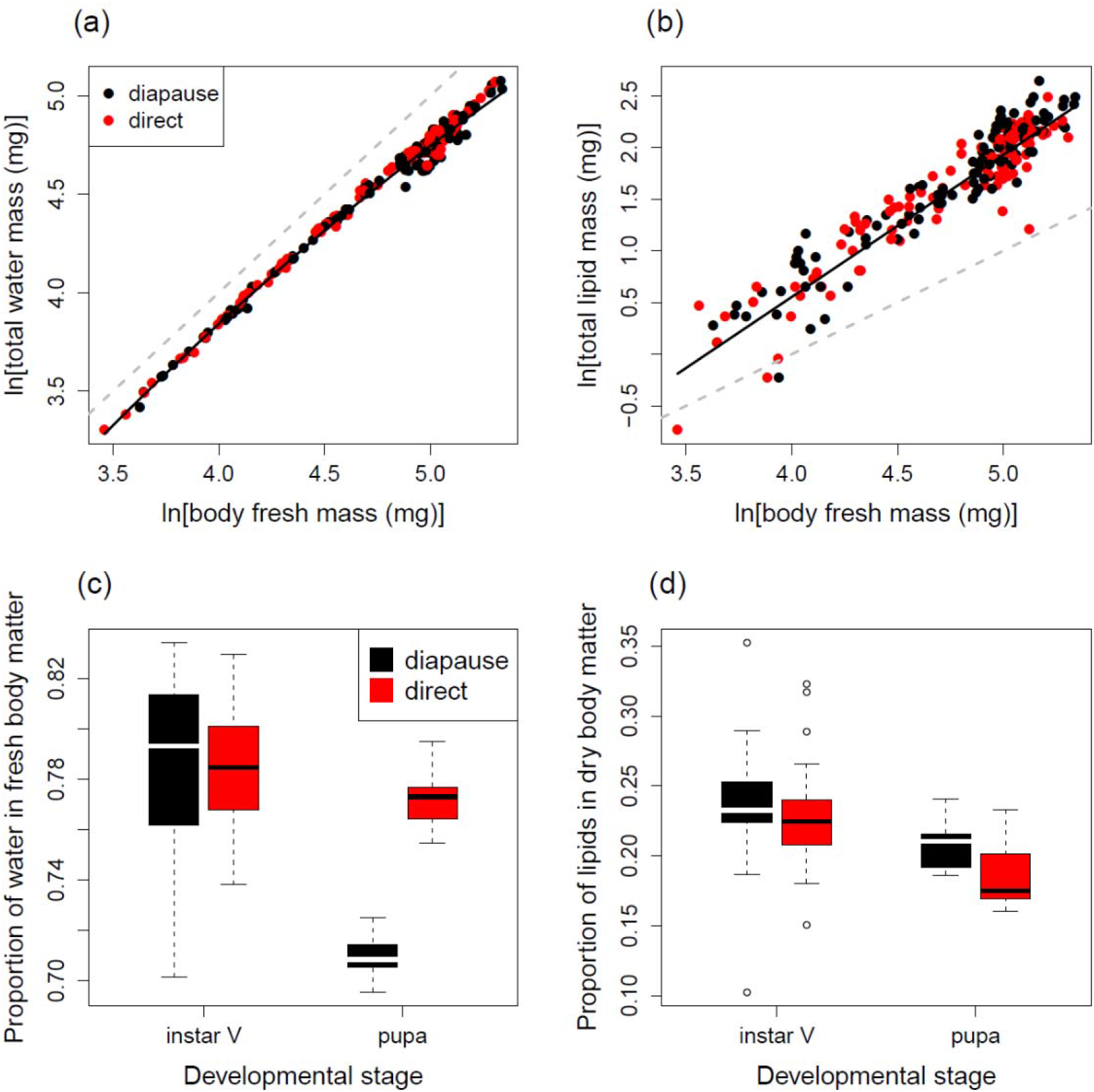
Instar V water content (a) and lipid content (b) in relation to body size in diapausing (closed circles) and directly developing (open circles) larvae, as well as comparison of proportion of water in fresh body matter (c) and proportion of lipids in dry body matter (d) between pupae and similarly-sized larvae. The unbroken curves in (a) and (b) are regression curves based on model-averaged coefficients (see Table 3), whereas the dashed grey lines are isometric reference lines with an arbitrary intercept chosen for the purpose of illustration. In (c) and (d), the boxes and whiskers illustrate the distribution of data similarly as in Fig. 1.

**Table 3.** Statistically significant (risk level 0.1; insignificant lower-level terms are presented for interactions and squared effects) model-averaged (full average) fixed effects of linear mixed-effects models fitted with the maximum likelihood method explaining variation in body water and lipid content between alternative developmental pathways (diapause / direct development), and developmental stages (instar V larva / pupa). The models included random family-specific intercepts.

| Trait | Term<br>(relative<br>importance) | Factor<br>levels | Averaged<br>estimate | Adjusted<br>S.E. | z-value | P-value |
| --- | --- | --- | --- | --- | --- | --- |
| Water<br>content V<sup>a</sup> |  | Intercept | -1.47 | 0.245 | 6.01 | <0.0001 |
|  | ln(fresh<br>mass) (1) |  | 1.67 | 0.113 | 14.7 | <0.0001 |
|  | [ln(fresh<br>mass)]<sup>2</sup> (1) |  | -0.0849 | 0.0130 | 6.55 | <0.0001 |
| Lipid<br>content V<sup>b</sup> |  | Intercept | -4.94 | 1.64 | 3.02 | 0.0026 |
|  | ln(fresh<br>mass)<br>(0.97) |  | 1.37 | 0.720 | 1.91 | 0.056 |
| Water<br>content:<br>larva vs.<br>pupa<sup>c</sup> |  | Intercept | 0.784 | 0.00604 | 130 | <0.0001 |
|  | Pathway<br>(1) | direct | 0.000393 | 0.00726 | 0.054 | 0.96 |
|  | Stage (1) | pupa | -0.0746 | 0.00642 | 11.6 | <0.0001 |
|  | pathway ×<br>stage (1) | direct ×<br>pupa | 0.0625 | 0.00805 | 7.76 | <0.0001 |
| Lipid<br>content:<br>larva vs.<br>pupa<sup>d</sup> |  | Intercept | 0.236 | 0.00646 | 36.7 | <0.0001 |
|  | Stage (1) | pupa | -0.0289 | 0.00888 | 3.25 | 0.0012 |

<sup>a</sup> ‘varExp’ variance function to take into account increasing residual variance with increasing fitted value.
<sup>b</sup> ‘varExp’ variance function to take into account decreasing residual variance with increasing fitted value.
<sup>c</sup> ‘varIdent’ variance function to take into account different residual variances in different combinations of developmental pathway and developmental stage (larva / pupa). <sup>d</sup> ‘varIdent’ variance function to take into account lower residual variance in pupae than in larvae.

When comparing two days old pupae with similarly-sized instar V larvae, relative water content was lower in diapause pupae than in larvae or directly developing pupae (Table 3; Fig. 5c). A similar comparison in the proportion of lipids in dry body matter revealed that the proportion of lipids was higher in instar V larvae than in pupae (Table 3; Fig. 5d), but there was no statistical support for a difference between the developmental pathways (Table 3).

## Discussion

We found the expected life history divergence between the diapause and direct development pathways through the last two larval instars in *P. napi;* larval growth and development were faster and body size smaller in individuals destined for direct development rather than pupal diapause. This life history differentiation between the alternative developmental pathways was associated with physiological differentiation, metabolic rate (defined as CO_2_ production) being higher under direct development than diapause. As expected, metabolic rate of larvae entering direct development and pupal diapause differed only in the final larval instar, and not in the penultimate instar. This is because the penultimate instar is the typical time window during which the developmental pathway is induced in *P. napi* (Friberg et al., 2011), resulting in a limited post-induction period for phenotypic divergence to become apparent in the penultimate instar. Putting these results together with the observation that the inter-pathway difference in development time became more pronounced in the final larval instar and the fact that the induction of a developmental pathway is largely independent of rates of growth and development in *P. napi* (Friberg et al., 2012; Kivelä et al., 2015), a high metabolic rate seems associated with fast growth and rapid development in individuals set for direct maturation after the induction of the developmental pathway. A high metabolic rate may be the cause for a high growth rate, or vice versa (Glazier, 2015). Either way, the relatively low metabolic rate in the diapause pathway and the lack of exaggerated lipid accumulation suggest that metabolic suppression is an important component of the diapause phenotype.

Ontogenetic allometric scaling of metabolism to body size was complex. The scaling of metabolic rate to body size was non-linear, early-instar hyperallometry turning into hypoallometry at the end of an instar (see also Glazier, 2005). This non-linear pattern was consistent both in the penultimate and final instars, yet the non-linearity was exaggerated in the penultimate instar. In an earlier study concentrating only on the diapause pathway, we found a similar non-linear scaling of metabolic rate to body mass in the penultimate instar in the diapause pathway and a linear scaling in the final instar (Kivelä et al., 2016b). A larger sample size in the present study is a likely explanation why we could detect a clearer levelling-off of metabolic rate in the penultimate instar (Figs 3a, c and S3) and a clear but weak non-linearity also in the final instar (Fig. 3b, d). Non-linearity of scaling may arise due to an increasing relative amount of metabolically inactive matter in the body towards the end of instars (see Glazier, 2005). Accordingly, the proportion of lipids in body matter largely follow this pattern. Moreover, the present data from the penultimate instar (see also Fig. S4) are consistent with the oxygen-dependent induction of moulting hypothesis, predicting that the tracheal respiratory system, that primarily grows during moults, reaches its maximum capacity at the end of the instar and so sets an upper limit for respiration rate (Greenlee and Harrison, 2004, 2005; Callier and Nijhout, 2011; Kivelä et al., 2016c).

Even though metabolic rate differed between the developmental pathways, there was no difference in the accumulation of lipids between diapause and direct development, although lipids constitute the main energy reserves used during diapause in many insects (Hahn and Denlinger, 2007, 2011; Arrese and Soulages, 2010), including *P. napi* (Lehmann et al., 2016). This is not exceptional (Hahn and Denlinger, 2007), yet unexpected given the ecology of *P. napi* where diapausing pupae must survive 5 to 10 months on the energy reserves acquired during the larval stage. In addition, many diapause-generation individuals pupate early in the summer and therefore also experience a long pre-winter warm period in pupal diapause. This may be particularly stressful as higher temperatures should lead to increased fat reserve depletion in ectotherms (Han and Bauce, 1998; Irwin and Lee, 2003; Bosch et al., 2010; Williams et al., 2014; see also Liu et al., 2016). Although there was no inter-pathway difference in the accumulation of energy reserves, final-instar *P. napi* larvae accumulate lots of lipids in their bodies during this instar, as indicated by the hyperallometric increase in total lipid mass in the body (Fig. 5b). Respiratory quotient (RQ) values exceeding one suggest that either metabolism is partially anaerobic (see e.g. Nielsen & Christian, 2007) or larvae convert carbohydrates to lipids (Zebe, 1953; Wigglesworth, 1972). Conversion of carbohydrates to lipids seems more likely as the proportion of lipids in dry body matter peaks similarly in halfway through the instar V as RQ (compare Fig. 4b to Fig. S5; see also Kivelä et al., 2016b). In the pupal stage, diapause pupae had lower water (and possibly higher lipid; see Fig. 4d) contents than directly developing ones soon after pupation and this difference emerged during pupation as larval water (and lipid) content did not differ between the developmental pathways. Reduced water content of diapause pupae may be a strategy to improve cold tolerance (see Danks, 2000; Bennett et al., 2005; Rozsypal et al., 2013).

Seasonal polyphenisms in life history and morphology associated with the diapause and direct development pathways have been demonstrated repeatedly in insects (Wiklund et al., 1991; Blanckenhorn and Fairbairn, 1995; West-Eberhard, 2003; Simpson et al., 2011; Aalberg Haugen et al., 2012; Friberg et al., 2012; Välimäki et al., 2013; Kivelä et al., 2013). Here, we show that the phenotypic divergence between the developmental pathways extends to metabolism, and the metabolic divergence appears after the induction of a developmental pathway has taken place. Pre-diapause metabolic divergence between larvae entering diapause and direct development that we demonstrate here persists after pupation (Lehmann et al., 2016). This is necessary because diapause pupae must survive up to 10 months with reserves similar to those of directly developing pupae that instead eclose as adults within a few weeks.

The generally extensive phenotypic divergence between alternative developmental pathways suggest that many components of the physiological and developmental machinery are independent enough for pathway-specific evolution. It is possible that pathway-specific selection changes metabolic rate, which would imply that metabolic rate is a key component of physiology affecting the evolution of life history polyphenisms (see also Arbačiauskas, 2004; Furness et al., 2015; but see Saastamoinen et al., 2013), and possibly life history evolution across species (Brown et al., 2004; Maino et al., 2014). Yet, it remains also possible that metabolic rate responds to changes in growth rate (Glazier, 2015), and that growth rate is the key to phenotypic differentiation between the alternative developmental pathways. Further studies are necessary for a rigorous assessment of these alternatives.

## Acknowledgements

We thank Loke von Schmalensee for help in running the experiment.

Author contributions
All authors conceived the study and designed the experiment. SMK and PL carried out the experiment. PL ran the body composition analyses and analysed the respirometric measurements. SMK ran statistical analyses. All authors participated in writing the manuscript.

