## Supplementary material for "Developmental plasticity in metabolism but not in energy reserve accumulation in a seasonally polyphenic butterfly"

**Electronic supplemental material**

^b^ Current address: Department of Ecology and Genetics, University of Oulu, PO Box 3000, 90014 University of Oulu, Finland

**Table S1.** Sets of models with Akaike weights >0.05 for the measured life history traits.

| **Variable** | **Model^a^** | **df** | **AIC_c_** | **ΔAIC_c_** | **Akaike weight** |
| --- | --- | --- | --- | --- | --- |
| Development time | 1/2/3 | 6 | 152.8 | 0 | 0.2 |
|  | 1/3 | 5 | 154.09 | 1.29 | 0.11 |
|  | 1/2/3/4 | 7 | 154.16 | 1.37 | 0.1 |
|  | 1/2/3/5 | 7 | 155.05 | 2.25 | 0.07 |
|  | 1/3/4 | 6 | 155.15 | 2.35 | 0.06 |
|  | 1/2/3/4/6 | 8 | 155.2 | 2.41 | 0.06 |
|  | 2/3 | 5 | 155.44 | 2.64 | 0.05 |
| Time until peak mass | 2/3 | 5 | 160.98 | 0 | 0.25 |
|  | 1/2/3 | 6 | 162.76 | 1.78 | 0.1 |
|  | 2/3/4 | 6 | 162.84 | 1.86 | 0.1 |
|  | 3 | 4 | 163.01 | 2.04 | 0.09 |
|  | 2/3/5 | 6 | 163.25 | 2.27 | 0.08 |
| Peak mass | 1/2/3/4 | 7 | 718.52 | 0 | 0.32 |
|  | 1/2/3/4/7 | 8 | 720.13 | 1.61 | 0.14 |
|  | 1/2/3 | 6 | 720.83 | 2.31 | 0.1 |
|  | 1/2/3/4/5 | 8 | 720.92 | 2.4 | 0.1 |
|  | 1/2/3/4/6 | 8 | 720.95 | 2.43 | 0.09 |
| Pupal mass | 1/2/3/4 | 7 | 683.18 | 0 | 0.2 |
|  | 1/3/4 | 6 | 684.36 | 1.19 | 0.11 |
|  | 2/3/4 | 6 | 684.36 | 1.19 | 0.11 |
|  | 1/2/3/4/7 | 8 | 685.48 | 2.3 | 0.06 |
|  | 1/2/3/4/6 | 8 | 685.55 | 2.37 | 0.06 |
|  | 3/4 | 5 | 685.56 | 2.39 | 0.06 |
|  | 1/2/3/4/5 | 8 | 685.61 | 2.43 | 0.06 |
| Growth rate IV^b^ | 2/3 | 10 | -132.84 | 0 | 0.25 |
|  | 2/3/5 | 11 | -130.86 | 1.98 | 0.09 |
|  | 1/2/3 | 11 | -130.80 | 2.05 | 0.09 |
|  | 3 | 9 | -130.73 | 2.11 | 0.09 |
|  | 2/3/4/6 | 12 | -130.30 | 2.54 | 0.07 |
|  | 2/3/4 | 11 | -130.25 | 2.60 | 0.07 |
| Growth rate V^c^ | 2/3/4/6/7 | 9 | -164.77 | 0 | 0.16 |
|  | 2/3/4/7 | 8 | -164.23 | 0.54 | 0.12 |
|  | 2/3 | 6 | -163.20 | 1.56 | 0.07 |
|  | 3/4/7 | 7 | -163.05 | 1.72 | 0.07 |
|  | 2/3/4/5/6/7/8 | 11 | -162.91 | 1.85 | 0.06 |
|  | 2/3/4/5/6/7 | 10 | -162.67 | 2.10 | 0.06 |

^a^ The numbers refer to model terms as follows: 1=cohort, 2=group, 3=pathway, 4=sex, 5=group×pathway, 6=group×sex, 7=pathway×sex; all models include random family-specific intercepts.

^b^ ‘varIdent’ variance function to take different variances in different families into account.

^c^ ‘varIdent’ variance function to take different variances in females and males into account.

**Table S2.** Sets of models with Akaike weights >0.05 for CO_2_ production rate in instars IV and V.

| **Variable** | **Model^a^** | **df** | **AIC_c_** | **ΔAIC_c_** | **Akaike weight** |
| --- | --- | --- | --- | --- | --- |
| CO2 IV^b^ | 1/2/3/4/5 | 10 | -127.32 | 0 | 0.15 |
|  | 1/2/4/5 | 9 | -126.82 | 0.5 | 0.12 |
|  | 1/2/3/4/5/6 | 11 | -125.17 | 2.15 | 0.05 |
| CO2 V^c^ | 1/2/3/4/5 | 11 | -262.99 | 0 | 0.15 |
|  | 1/2/4/5 | 10 | -262.1 | 0.89 | 0.1 |
|  | 1/2/3/4/5/9 | 12 | -261.48 | 1.51 | 0.07 |
|  | 1/2/3/4/5/6 | 12 | -261.07 | 1.93 | 0.06 |
|  | 1/2/3/4/5/7 | 12 | -261.03 | 1.97 | 0.06 |

^a^ The numbers refer to model terms as follows: 1=ln(centered mass), 2= ln(centered mass)^2^, 3=cohort, 4=frass, 5=pathway, 6=sex, 7= ln(centered mass)×pathway, 8= ln(centered mass)×sex,

9= ln(centered mass)^2^×pathway

^b^ Random individual-specific intercepts and slopes in relation to centered mass.

^c^ Random individual-specific intercepts and slopes in relation to centered mass; ‘varIdent’ variance function to take into account higher residual variance in females than males.

**Table S3.** Sets of models with Akaike weights >0.05 for respiratory quotient (RQ) in instars IV and V.

| **Variable** | **Model^a^** | **df** | **AIC_c_** | **ΔAIC_c_** | **Akaike weight** |
| --- | --- | --- | --- | --- | --- |
| RQ IV | 1 | 5 | -29.01 | 0 | 0.07 |
|  | 1/3/4 | 7 | -28.94 | 0.07 | 0.07 |
| RQ V | 1/3/4 | 6 | -419.32 | 0 | 0.12 |
|  | 1/2/3/4 | 7 | -418.94 | 0.38 | 0.10 |
|  | 1/3/4/6 | 7 | -418.32 | 1.00 | 0.07 |
|  | 1/2/3/4/6 | 8 | -418.11 | 1.21 | 0.06 |

^a^ The numbers refer to model terms as follows: 1=cohort, 2=frass, 3=mass,

4=mass^2^, 5=pathway, 6=sex

**Table S4.** Sets of models with Akaike weights >0.05 for water and lipid content in instar V.

| **Variable** | **Model^a^** | **df** | **AIC_c_** | **ΔAIC_c_** | **Akaike weight** |
| --- | --- | --- | --- | --- | --- |
| Water content V^b^ | 1/2/3/4/5 | 9 | -855.99 | 0 | 0.32 |
|  | 1/2/3 | 7 | -855.52 | 0.47 | 0.25 |
|  | 1/2/3/5 | 8 | -855.18 | 0.81 | 0.21 |
|  | 1/2/3/4 | 8 | -854.96 | 1.03 | 0.19 |
| Lipid content V^c^ | 1/3/4 | 7 | -68.15 | 0 | 0.27 |
|  | 13 | 6 | -67.64 | 0.51 | 0.21 |
|  | 1/2/3/5 | 8 | -66.60 | 1.55 | 0.13 |
|  | 1/2/3/4/5 | 9 | -66.54 | 1.61 | 0.12 |
|  | 1/2/3/4 | 8 | -66.37 | 1.78 | 0.11 |
|  | 1/2/3 | 7 | -65.93 | 2.22 | 0.09 |

^a^ The numbers refer to model terms as follows: 1=ln(fresh mass), 2= [ln(fresh mass)]^2^, 3=pathway, 4=ln(fresh mass)×pathway, 5= [ln(fresh mass)]^2^×pathway; all models included random family-specific intercepts.

^b^ ‘varExp’ variance function to take into account increasing residual variance with increasing fitted value.

^b^ ‘varExp’ variance function to take into account decreasing residual variance with increasing fitted value.

**Table S5.** Sets of models with Akaike weights >0.05 for the comparison of relative water and lipid content in pupae and similarly-sized instar V larvae.

| **Variable** | **Model^a^** | **df** | **AIC_c_** | **ΔAIC_c_** | **Akaike weight** |
| --- | --- | --- | --- | --- | --- |
| Water content: larva vs. pupa^b^ | 1/2/3 | 9 | -497.33 | 0 | 1 |
| Lipid content: larva vs. pupa^c^ | 1/2/3 | 7 | -435.53 | 0 | 0.56 |
|  | 1/2 | 6 | -435.01 | 0.52 | 0.43 |

^a^ The numbers refer to model terms as follows: 1=pathway, 2=(developmental stage), 3=pathway×(developmental stage); all models included random family-specific intercepts.

^b^ ‘varIdent’ variance function to take into account different residual variances in different combinations of developmental pathway and developmental stage (larva / pupa).

^c^ ‘varIdent’ variance function to take into account lower residual variance in pupae than in larvae.

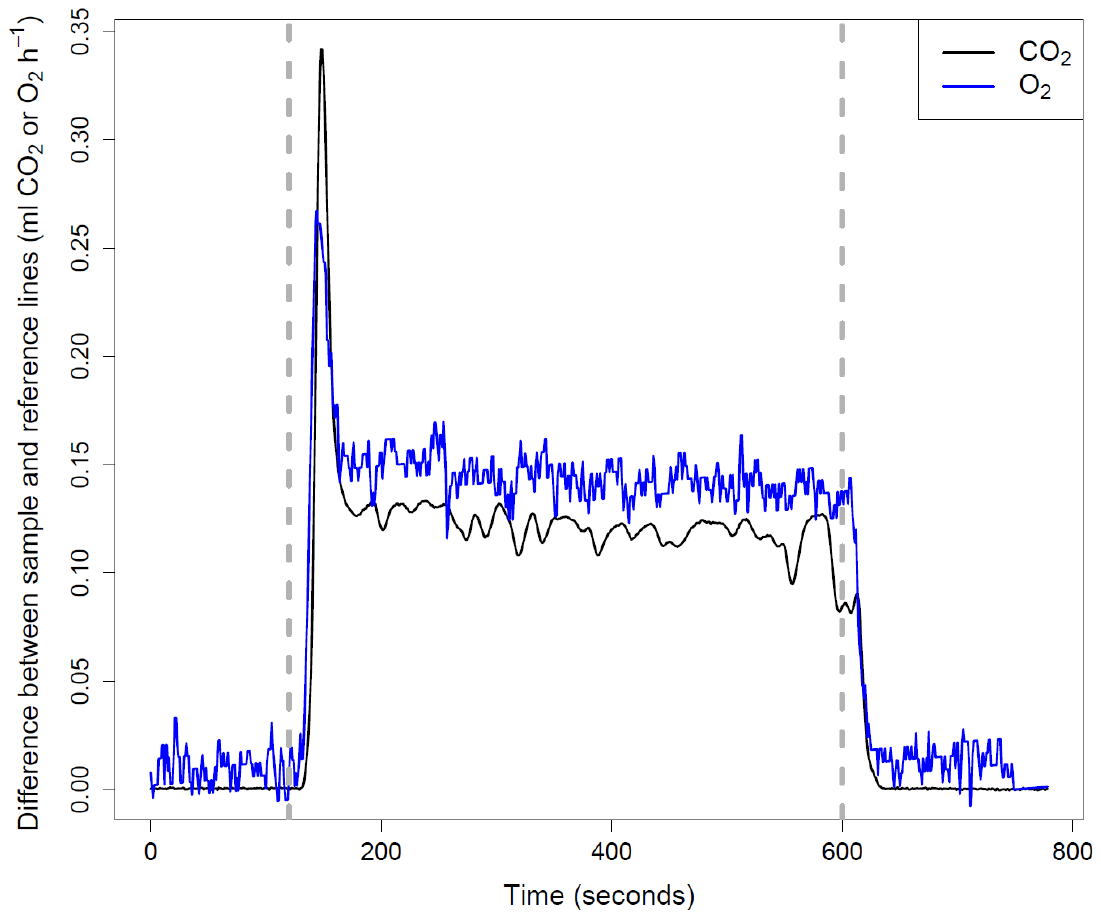

**Figure S1.** **An example of CO_2_ (black line) and O_2_ (blue line) traces during a respirometric measurement.** The vertical grey dashed lines indicate the start and end of the measurement of the sample line. The O_2_ signal is lag-corrected in relation to CO_2_ signal. This particular example shows a measurement of an instar V larva weighing 162.2 mg.

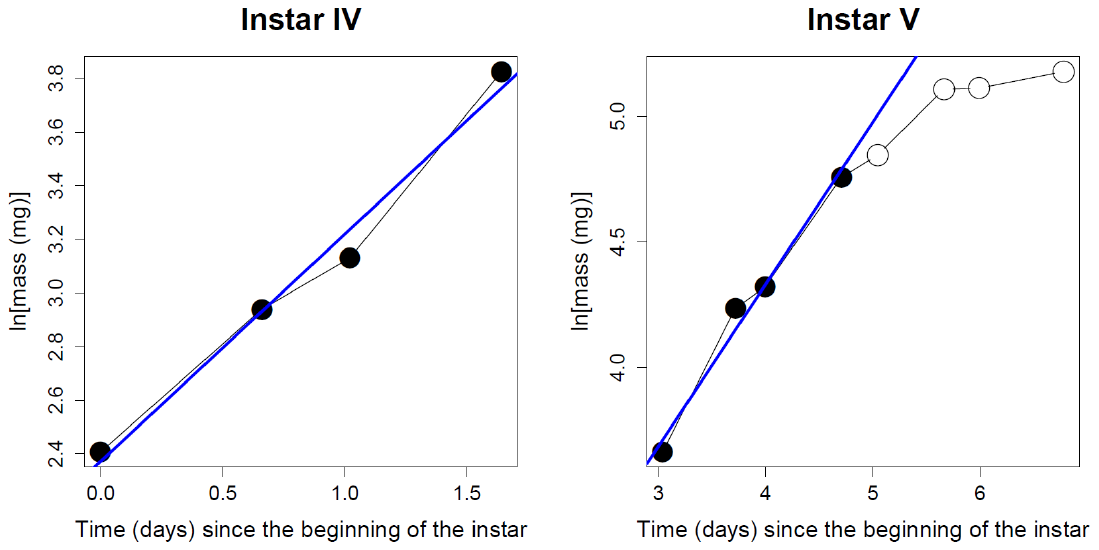

**Figure S2.** **Illustration of growth rate calculation.** Individual-specific growth rate estimate in the instar IV (left) and V (right) is the slope of the regression line (blue lines in the figure) of ln(mass) on time (days) in the focal instar. The figure illustrates the observed growth trajectory of a single individual (circles connected with black lines) through the two instars. Only growth period data are shown (i.e. pre-moult mass-loss is excluded in both instars). Only observations marked with closed circles were included in the regression-based estimation of growth rate. All observations were included in instar IV, whereas only the first 50% of observations (rounded to an integer number of observations) were included in instar V to exclude growth deceleration due to preparation to pupation from estimation. Therefore, the growth rate estimates pertain to period of free growth.

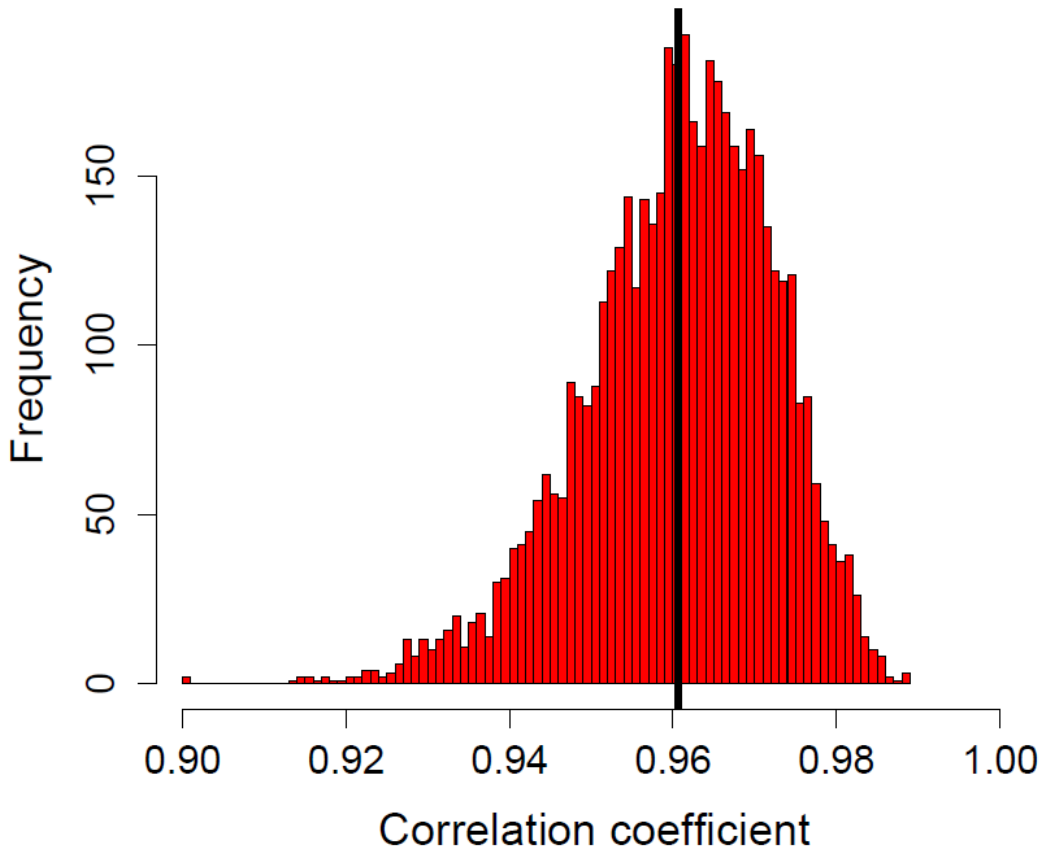

**Figure S3.** **Distribution of correlation coefficients between CO_2_ production rate and metabolic rate that is based on resampling.** To avoid problems due to non-independency of within-individual measurements, 5000 resamples were derived so that each individual was represented by a single randomly-drawn observation in each round, and correlation coefficient (Pearson correlation) between CO_2_ production rate and metabolic rate was calculated from each of the 5000 resamples. The vertical line indicates the mean of the distribution. Metabolic rate (MR) was calculated from respirometry data as MR = (*X* × V̇O_2_) / 3600, where *X* = 15.97 + (5.164 × RQ) is the oxyjoule equivalent (J [ml O_2_]^-1^), and RQ is the respiratory quotient (V̇CO_2_ / V̇O_2_; V̇CO_2_ is CO_2_ production rate [ml CO_2_] h^-1^; V̇O_2_ is O_2_ consumption rate [ml O_2_] h^-1^) (see Lighton et al. 1987). The denominator, 3600 s (1 h = 3600 s), converts MR into Watts (i.e. J s^-1^).

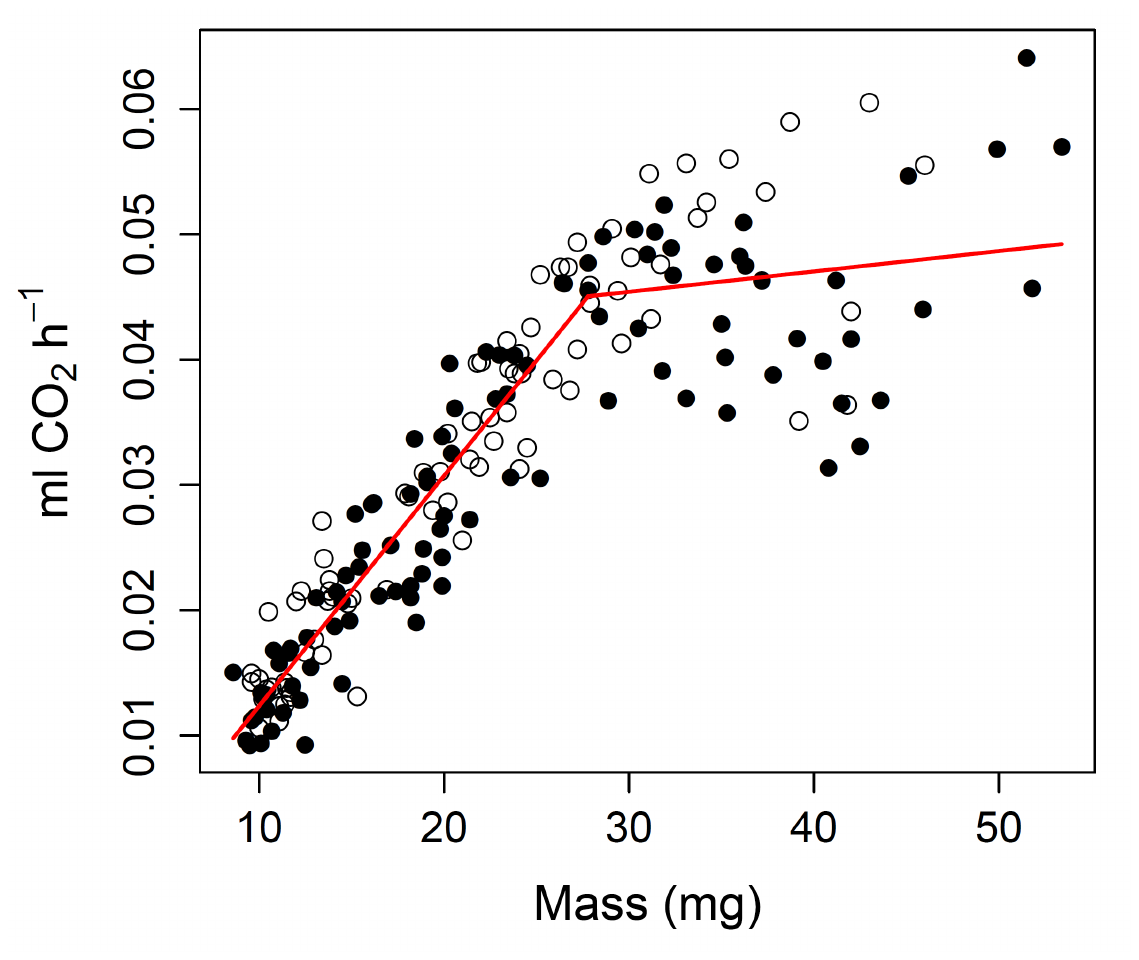

**Figure S4.** **Untransformed CO_2_ production in the penultimate larval instar in relation to body mass under diapause (closed circles) and direct development (open circles).** The red line is a fitted regression line from a segmented regression analysis (Muggeo 2003), where a breakpoint in the slope is estimated. The segmented regression analysis was run with the R package ‘segmented’ (Muggeo 2008). The estimated breakpoint that can be considered to be a surrogate of the so-called critical mass for moult induction (see Callier and Nijhout 2011) is 27.8 (95% CI: 26.0, 29.6) mg, the regression slope being 0.00184 (0.00167, 0.00201) below the breakpoint and 0.000162 (-4.19×10^-5^, 0.000367) above it.

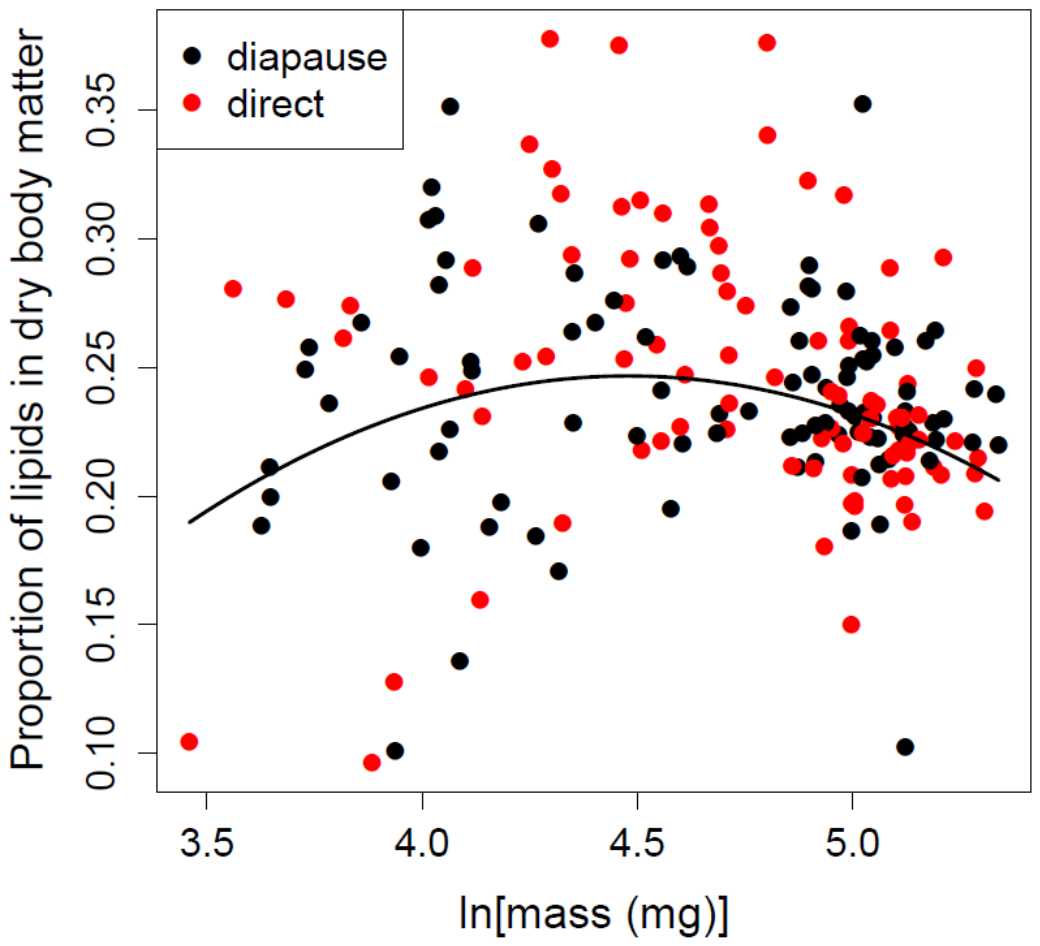

**Figure S5. Proportion of lipids in dry body matter in instar V in relation to body size in larvae entering pupal diapause (black symbols) and direct development (red symbols).**
